# Thalamocortical boutons cluster by ON/OFF responses in mouse primary visual cortex

**DOI:** 10.1101/2022.09.26.509569

**Authors:** Elaine Tring, Dario L. Ringach

**Affiliations:** Department of Neurobiology, David Geffen School of Medicine, UCLA, Los Angeles, CA 90095; Department of Psychology, UCLA, Los Angeles, CA 90095

**Keywords:** ON/OFF domains, thalamocortical projection, biased input, canonical correlation, mouse

## Abstract

In higher mammals, the thalamic afferents to primary visual cortex cluster according to their responses to increases (ON) or decreases (OFF) in luminance. This feature of thalamocortical wiring is thought to create columnar, ON/OFF domains in V1. We have recently shown that mice also have ON/OFF cortical domains, but the organization of their thalamic afferents remains unknown. Here we measured the visual responses of thalamocortical boutons using two-photon imaging and found they also cluster in space according to ON/OFF responses. Moreover, fluctuations in the relative density of ON/OFF boutons mirrors fluctuations in the relative density of ON/OFF receptive field positions on the visual field. These findings indicate that there is a segregation of ON/OFF signals already present in the thalamic input, a result that may reflect the spatial organization of retinal ganglion cell mosaics themselves.

**New and Noteworthy:** Neurons in primary visual cortex cluster into ON and OFF domains, which have shown to be linked to the organization of receptive fields and cortical maps. Here we show that in the mouse, such clustering is already present in the thalamic input, suggesting that the cortical architecture is shaped by the periphery.

## Introduction

In higher mammalian species, geniculate afferents cluster according to their ON/OFF responses (1–4). This feature of thalamocortical wiring is widely believed to induce columnar, ON/OFF cortical domains, where the responses of neurons are dominated by either the onset (ON) or offset (OFF) of luminance within their receptive fields (2–6). We have recently found that the primary visual cortex in the mouse is also parceled into ON/OFF domains. If the origin of ON/OFF cortical domains is a consequence of the clustering of afferents by ON/OFF type, one would predict ON/OFF thalamic inputs to segregate in the mouse as well. Indeed, our main finding is that thalamocortical boutons in the mouse are spatially clustered according to their responses to light increments and decrements. Moreover, fluctuations in the dominance of ON/OFF responses in thalamic boutons mirror fluctuations in density of ON/OFF receptive fields across the visual field. This latter finding suggests that spatial biases in the representation of ON and OFF responses are likely to reflect a property of retinal ganglion cell mosaics.

Understanding the organization and origin of ON/OFF domains has been attracting increased attention because their spatial layout is linked to the structure of receptive fields and cortical maps, including those for orientation, direction, and retinal disparity (2, 6–16). Simple cells tend to be located between the centers of ON and OFF domains, with their receptive fields organized into ON/OFF subregions matching the retinotopic location of ON/OFF signals in nearby domains (6), in both species with and without orientation maps (17). Thus, the question of how ON/OFF domains are formed is of outmost relevance to our understanding of how the cortical architecture is established. If, as we propose, the origin of ON/OFF domains can be traced back to the statistics of the retinal ganglion cell mosaics in the retina, the results would support the notion that the wiring of receptive fields and the organization of the cortical architecture is initially seeded by the structure of signals from the periphery (11, 12, 14, 18–20). The present results lend support this hypothesis by showing that the clustering of boutons in V1 is not due to a spatial selection of the afferents by type by cortical sites, but rather their spatial structure mimics fluctuations in the representation of ON/OFF signals already present in the visual field.

## Materials and Methods

### Animals

All procedures were approved by UCLA’s Office of Animal Research Oversight (the Institutional Animal Care and Use Committee) and were in accord with guidelines set by the U.S. National Institutes of Health. A total of 4 C57BL/6J WT mice, male (1) and female (3), aged P35-56, were used in the study. Data from six different cortical volumes were obtained.

### Surgery

Two-photon imaging experiments were conducted through chronically implanted cranial windows over primary visual cortex. Carprofen was administered pre-operatively (5mg/kg, 0.2mL after 1:100 dilution). Mice were anesthetized with isoflurane (4%–5% induction; 1.5%–2% surgery) and body temperature was maintained at 37.5C using a feedback heating system. Eyes were coated with a thin layer of ophthalmic ointment to prevent desiccation. Anesthetized mice were mounted in a stereotaxic apparatus using blunt ear bars in the external auditory meatus to immobilize the head. A portion of the scalp overlying the two hemispheres of the cortex (approximately 8mm by 6mm) was removed to expose the underlying skull. The skull was dried and covered by a thin layer of Vetbond. After a drying period (15 min) the Vetbond provided a stable and solid surface to affix an aluminum bracket (a head holder) with dental acrylic. The bracket was then affixed to the skull and the margins sealed with Vetbond and dental acrylic to prevent infections. A craniotomy over monocular V1 on the left hemisphere was conducted using a high-speed dental drill. Special care was taken to ensure that the dura was not damaged during the process. A stock concentration of AAV1-CAG-GCaMP6S (Addgene: AAV1.CAG.GCaMP6s.WPRE.SV40; #100844-AAV1; titer: ∼2e13 GC/ml) was pressure injected using a PicoSpritzer III (Parker, Hollis, NH). We used a thin-walled glass pipette (Warner Instruments, #64-0800) pulled by Sutter P-1000 to create a sharp injection pipette (∼0.3-0.7 μm tip), then the last 1-2 mm of the tip is broken to create a 6 μm tip for injection. The injection pipette was filled with the virus and positioned over the dorsal LGN with coordinates 2.1 mm posterior from bregma and 2.3 mm lateral from the midline using a micromanipulator. Then, the pipette was slowly lowered to a depth of 2800 μm below the pial surface. Starting at a depth of 2800 μm, 10 puffs were given at 15-20 psi with a duration of 10 ms, each puff is separated by a 4s interval and making injections every 10 μm moving up, with the last injection made at a depth of 2600 μm. The total volume injected was approximately 0.2 μl. A sterile 3 mm diameter cover glass was placed directly on the exposed dura and sealed to the surrounding skull with Vetbond. The remainder of the exposed skull and the margins of the cover glass were sealed with dental acrylic. Mice were allowed to recover on a heating pad. Once alert, mice were transferred back to their home cage. Carprofen was administered post-operatively for 72 hours. Mice were allowed to recover from surgery for two weeks, after which we began monitoring the levels of expression of GCaMP6s in thalamic boutons in V1. If the expression levels were found to be adequate, we moved on to measure the visual responses of boutons and analyze their distribution in the cortex as detailed below.

### Two-photon imaging

At the beginning of each session mice were briefly sedated and administered Texas red (ThermoFisher #D3328, 0.1 mL, s.c, from a 2 mg/mL in PBS stock). Mice were positioned on a running wheel and head-restrained under a resonant, two-photon microscope (Neurolabware, Los Angeles, CA) controlled by Scanbox acquisition software and electronics (Scanbox, Los Angeles, CA). After a waiting period of ∼20 min, we were able to image both geniculate boutons and the cortical vasculature on the green and red PMT channels respectively (**Fig 1A**). The light source was a Coherent Chameleon Ultra II laser (Coherent Inc, Santa Clara, CA). Excitation wavelength was set to 920nm. The objective was an x16 water immersion lens (Nikon, 0.8NA, 3mm W.D.). The microscope frame rate was 15.6Hz (512 lines with a resonant mirror at 8kHz). The field of view was 516*µ*m × 308*µ*m in all instances. The objective was tilted to be approximately normal the cortical surface. An electronically tuned lens (Optotune EL-10-30-C, Dietikon, Switzerland) was used to run independent sessions acquiring data from optical planes spaced 15*µ*m apart (to ensure disjoint sets of boutons in different sections) starting at a depth of ∼250*µ*m from the cortical surface (**Fig 1B**). A total of 6 datasets from 4 mice were recorded each with different number of optical sections (see table below). Images were processed using a standard pipeline consisting of image registration (based on the Texas red signal), cell segmentation and deconvolution using Suite2p (https://suite2p.readthedocs.io/). For any one optical section, the location of the cells in the imaging plane were estimated as the center of mass of the corresponding region of interest calculated by Suite2p.

**Fig 1.**
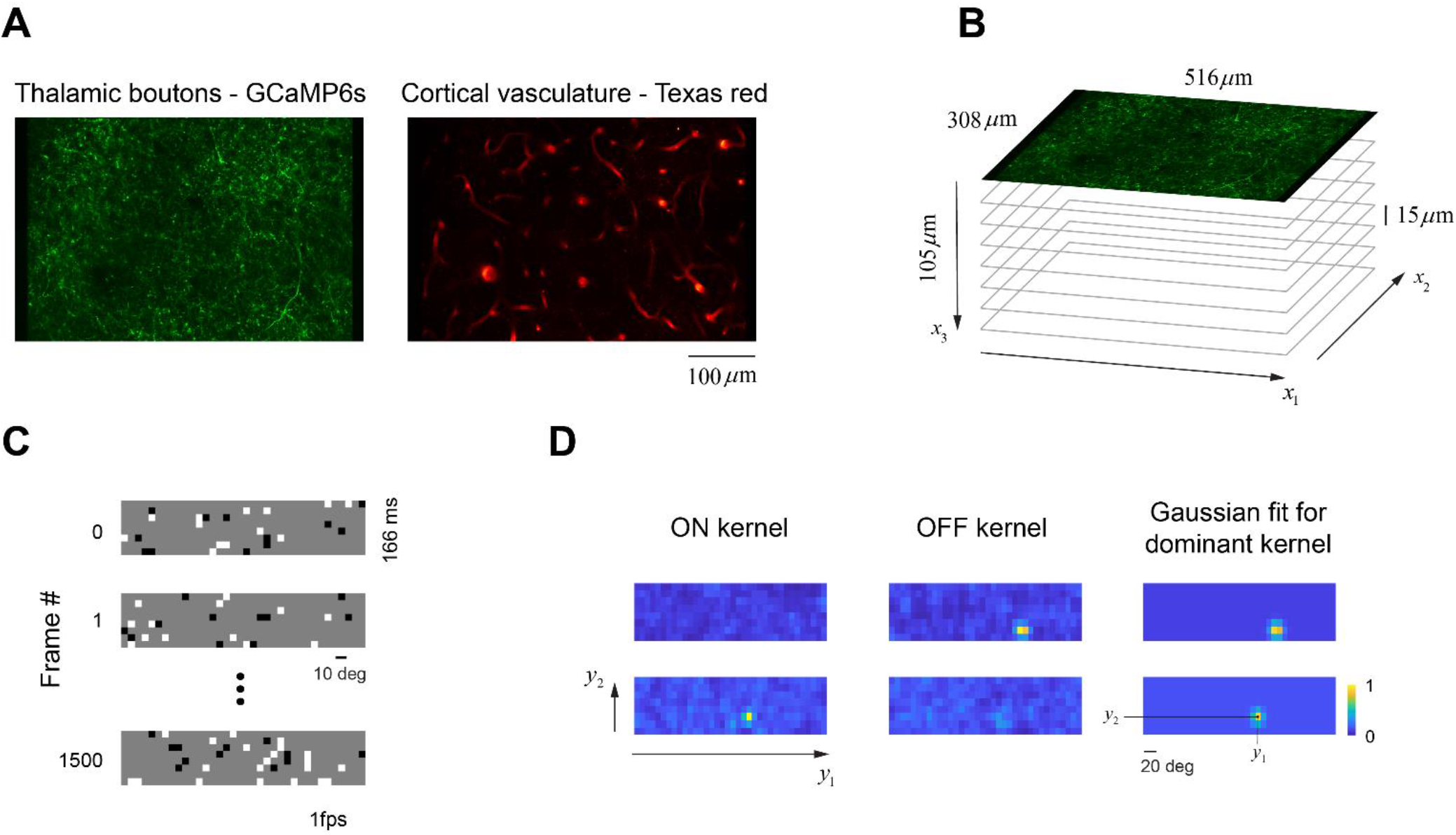
Two-photon imaging from thalamic boutons in V1. (**A**) Green PMT channel showing the expression of GCaMP6s in thalamic afferents and boutons. The red PMT channel shows a visualization of the vasculature by injections of Dextran Texas red, which assisted in the registration of the images during our analyses. **(B)** A typical experiment consisted in sampling a volume in V1 where adjacent optical sections were separated by 15*µ*m. **(C)** Sparse noise stimulation was used to evoke response from thalamic boutons. **(D)** Reverse correlation of signals from each individual bouton with bright and dark stimuli led to ON and OFF kernels in each case. Shown are two examples of a bouton responding largely to bright stimuli (ON bouton) and one responding largely to dark stimuli (OFF bouton). The panels on the right illustrate two-dimensional Gaussian fits to the dominant kernels, which yield an estimate of the location of the bouton’s receptive field in visual space.

A camera synchronized to the frame rate of the microscope imaged the contralateral eye during data collection. These data were subsequently analyzed to determine the center and size of the pupil within the image plane. The distribution of eye movements was computed, yielding a mode and standard deviation. There were no obvious differences between the analyses performed on the entire dataset or on data segments where the eye position was restricted to lie within 1SD of the mode. Here, we report the analysis using the entire dataset.

A summary of the datasets available is provided by the table below:

Across all the experiments, ON boutons were the most numerous representing 68% of the population, while OFF boutons accounted for 26% of the population (**Table 1**). About 6% of boutons showed both ON and OFF kernels and were not considered any further in our analyses.

**Table 1.** Summary of experiments. Each line summarizes each of the experiments conducted including the sex of each animal, the number of optical sections (planes) obtained, the total number of ON and OFF boutons that pass the data selection criteria, the number of boutons with both ON and OFF response, and the correlation and p-value between the fluctuations the spatial distribution of ON/OFF responses in boutons on the cortex and the fluctuations of the ON/OFF receptive fields centers on the visual field. The shaded lines correspond to the sample data shown in Figures 2 and 3.

| Dataset # | mouse ID | sex | # planes | $N_{on}$ | $N_{off}$ | $N_{on+off}$ | Correlation | p-value |
| --- | --- | --- | --- | --- | --- | --- | --- | --- |
| 1 | LGNP01 | F | 6 | 631 | 139 | 28 | 0.51 | 0.003 |
| 2 | LGNP07 | M | 7 | 248 | 115 | 38 | 0.30 | 0.045 |
| 3 | LGNP07 | M | 6 | 519 | 351 | 315 | 0.64 | 0.001 |
| 4 | LGNP11 | F | 4 | 258 | 88 | 20 | 0.34 | 0.050 |
| 5 | LGNP11 | F | 7 | 381 | 186 | 23 | 0.43 | 0.005 |
| 6 | LGNP12 | F | 10 | 191 | 59 | 4 | 0.43 | 0.012 |

### Visual stimulation

A Samsung CHG90 monitor positioned 30 cm in front of the animal was used for visual stimulation. The screen was calibrated using a Spectrascan PR-655 spectro-radiometer (Jadak, Syracuse, NY), generating gamma corrections for the red, green, and blue components via a GeForce RTX 2080 Ti graphics card. Visual stimuli were generated by a Processing sketch written by our laboratory using OpenGL shaders (see http://processing.org). The screen was divided into an 18 by 8 grid, resulting in an tile size of 8 by 8 deg, thereby matching the size of a typical LGN center (21) (**Fig 1C**). Each frame of the stimulus was generated by selecting the luminance of each tile randomly as either bright (10% chance), dark (10% chance) or grey (80% chance), with 100% contrast. The stimulus was flashed for 166 ms and appeared at a rate of 1 per second. The screen was uniform gray between stimuli. Transistor-transistor logic (TTL) pulses generated by the stimulus computer signaled the onset of stimulation. These pulses were time-stamped by the microscope with the frame and line number that was being scanned at that moment the signals occurred. Sessions lasted for 25 min, generating the response of cells in the population to 1500 stimulus presentations.

### Calculation of ON and OFF kernels

For each bouton and tile in the stimulus we calculated the average response the bouton locked to the presentation of bright or dark stimulus over the 15 frames (1 sec) following stimulus onset. The ON kernel at a delay of *t* frames after stimulus onset is represented as an image of equal size to the stimulus. The value of this image at tile location (*i, j*) corresponds to the average response following the presentation of a bright stimulus at that location *t* frames after stimulus onset. We denote this image by *ON*(*t*) and adopt a similar definition of the OFF kernel, *OFF*(*t*). For each time delay, we compute the norm of the kernel normalized by the norm at *t* = 0: *S*_*on*_(*t*) = ‖*ON*(*t*)‖/‖*ON*(0)‖ and, similarly we calculate *S*_*off*_(*t*) = ‖*OFF*(*t*)‖/‖*OFF*(0)‖. These curves typically peak at delays of ∼5 frames (corresponding to ∼320ms). We declared a bouton to have a significant ON kernel if its normalized norm attained a peak value larger than 5 and a two-dimensional Gaussian fit of the kernel at the peak delay time accounts for at least 50% of its variance. A similar definition applied to OFF kernels. As a result, a bouton could have no significant maps, either significant ON or OFF maps, or both (see **Table 1**). The two-dimensional Gaussians fits to the dominant ON and OFF kernels yield their center locations (*y*_1_, *y*_2_) on the visual field (**Fig. 1D**).

### Canonical correlation analysis

Each bouton was assigned a coordinate in cortical space, (*x*_1_, *x*_2_, *x*_3_) (**Fig 1B**) and, for its dominant kernel, one in visual space, (*y*_1_, *y*_2_) (**Fig 1D**). Canonical correlation analysis finds transformations 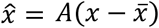 and 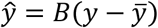 such that the covariance of 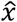 and *ŷ* is diagonal and the correlations between matching canonical coordinates are maximized. The transformations are further constrained so that the variance of the canonical coordinates equals one. Note that in our case the matrix *A* is *n* × 3, while the matrix *B* is *n* × 2, where *n* is the total number of cells with at least one significant map. We used MATLAB’s `cannoncorr()` function for these analyses. The goal of canonical correlation analysis to represent cortical and visual space on a common two-dimensional latent space. The result is an optimal alignment of cortical and visual space under an affine transformation.

### Density estimation

Given a distribution of points in native or canonical space (either cortical or visual) we estimate the density distribution by 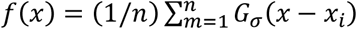, where *G*_*σ*_(·) is a two-dimensional Gaussian kernel of width *σ* and {*x*_*i*_} (*i* = 1, …, *N*) is the set of points under consideration (22). For canonical variables, we chose a width of *σ* = 0.25, following the rule of thumb bandwidth estimator 0.9 *n*^−1/5^, with *n*∼500, which is a typical size for our data (22). In native cortical space, we used *σ* = 30μ*m*. Estimates of 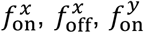 and 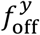 and their counterparts in canonical space were all obtained by this procedure.

To evaluate the likelihood that the observed fluctuations could arise by chance, we randomly shuffled the labels of ON and OFF cells in *N* = 1000 experiments. For each experiment, we calculated the distribution of fluctuations at each point in cortical and visual spaces, which enabled us to compute *p* = 0.001 level sets. Similarly, we computed the distribution of correlation coefficients between fluctuations of ON/OFF cells on the cortical surface and those of their center locations in the visual field, allowing us to calculate the statistical significance of the observed correlation in the original data.

### Data and code availability

The data from all experimental sessions, including the processed kernels and estimated parameters, have been deposited at _________________. Sample code describing the structure of the database and the calculation of ON/OFF bouton distributions is provided along with of the data in the same Figshare repository.

## Results

We used two-photon imaging in alert, head-fixed mice to measure the responses of thalamic boutons in a volume of primary visual cortex to visual stimuli. We expressed GCaMP6s in the LGN by injections of pAAV.CAG.GCaMP6s.WPRE.SV40 (**Fig 1A**, left panel) and characterized the responses of individual boutons using two-photon imaging. To assist in the registration of our images, we labeled the vasculature with Texas red (**Fig 1A**, right panel). We used sparse-noise stimulation (**Fig 1C**) to record the responses of thalamocortical bouton within a cortical volume sampled with 3 − 6 optical sections spaced 15 *µ*m apart (**Fig 1B**). A standard data analysis pipeline comprised of image registration (based on the red PMT channel), cell segmentation, signal extraction and deconvolution steps, yielded the estimated spiking of synaptic boutons. The centroid of the regions of interest (ROIs), along with the depth of the optical section, allowed us to assign each bouton a coordinate in cortical space, (*x*_1_, *x*_2_, *x*_3_) (**Fig 1B**). As the objective was nearly normal to the cortex, (*x*_1_, *x*_2_) represents the projection of a bouton on the cortical surface and *x*_3_ represents its depth.

We computed the ON and OFF receptive fields (or kernels) of each bouton in the population by correlating their responses with the locations of bright and dark patches in the stimuli at different time delays (**Fig 1D**). We defined ON/OFF kernels at the optimal delay for which the norm of the kernel attained its maximum norm. Boutons that had only a statistically significant ON kernel (68% of the population) were defined as ON (**Fig 1D**); a similar definition applied to OFF (26% of the population). A small fraction of boutons (∼6% of the population) showed both ON and OFF kernels and were not consider further in the analysis.

### Thalamocortical boutons cluster by ON/OFF types

To test if boutons cluster in cortical space according to their type we computed the difference in the spatial density of ON and OFF boutons. Given a set of points 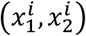 corresponding to ON boutons (ignoring their depth) we estimated their density via a kernel estimate, 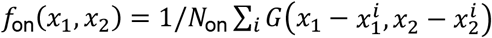, where *N*_on_ is the number of ON boutons and *G*(·) is a Gaussian kernel(22) (Methods). A similar density estimate can be obtained for OFF boutons, as 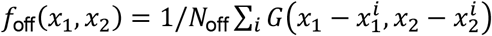. The fluctuation in the density of ON and OFF boutons is given by the difference *f*_on_(*x*_1_, *x*_2_) − *f*_off_(*x*_1_, *x*_2_). We observed that *f*_on_(*x*_1_, *x*_2_) and *f*_off_(*x*_1_, *x*_2_) have non-uniform distributions that peak at different locations (**Fig. 2**). As a result, the difference *f*_on_(*x*_1_, *x*_2_) − *f*_off_(*x*_1_, *x*_2_) showed a clear spatial structure with regions where the density of ON cells is higher than OFF cells (ON domains), and regions where the density of OFF cells was higher than ON cells (OFF domains). The statistical significance of these fluctuations was assessed by Monte Carlo simulations where the ON and OFF labels of the cells are randomly shuffled, allowing us to determine the locations where the deviations attained a significance at the 0.001 level (**Fig. 2**, right panels, blue and red level sets). These results are typical of our datasets, all of which showed regions with statistically significant ON and OFF dominance. We conclude that thalamocortical boutons in V1 are organized into ON/OFF domains where responses of one type dominate over the other. We emphasize the segregation is far from perfect, but the analysis certainly demonstrates that boutons cluster by ON/OFF type beyond what might be expected by chance.

**Fig 2.**
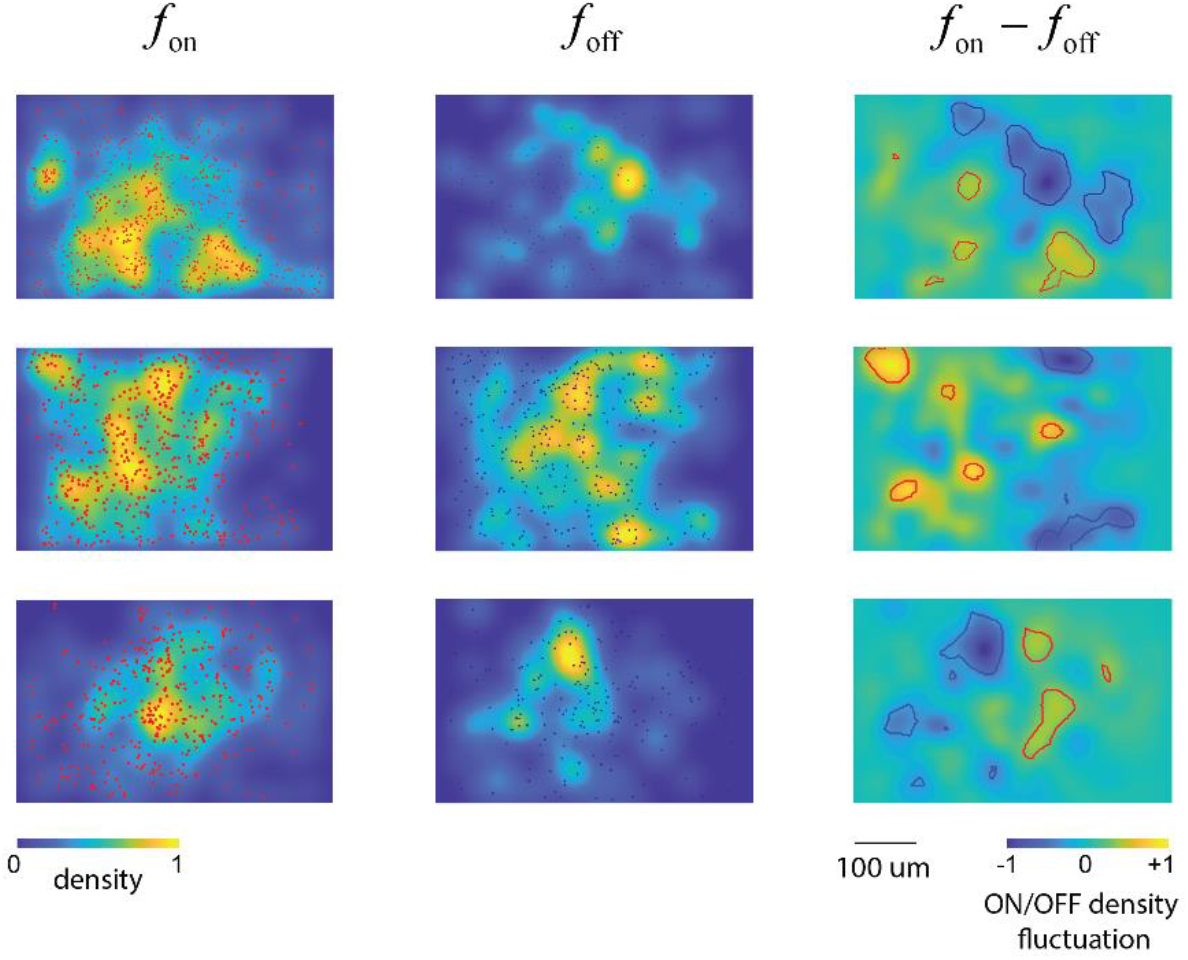
Clustering of ON/OFF boutons. Each row corresponds to a separate experiment. The panel on the left shows the distribution of ON boutons (projected onto the cortical surface by disregarding their depth). The middle panel shows the density of OFF boutons. The panel on the right, shows the fluctuations in density. Red and blue contours represent areas where the difference in density exceeds the magnitude of what would be expected by randomly shuffling the ON/OFF levels of the boutons at a significance level of p=0.001.

### Fluctuations in ON/OFF boutons correlates with fluctuations of ON/OFF receptive fields centers

A parsimonious explanation for the clustering of ON/OFF boutons is that there might already be an imbalance in the density of ON/OFF center receptive fields on the visual field, originating in the retinal ganglion cell mosaics, that is maintained due to retinotopic mapping in the thalamus and cortex. This hypothesis predicts a correlation between the fluctuations between ON/OFF signals in thalamic boutons and fluctuations in the density of their receptive fields on the visual field (17).

To test this prediction we bring cortical and visual fields into alignment using canonical correlation analysis (23), which generates two linear transformations of the data. Cortical space is mapped to 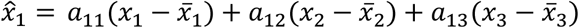 and 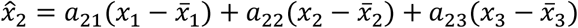, or in matrix form 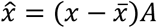. Similarly, the visual field is mapped by 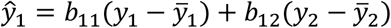 and 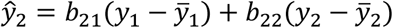, or in matrix form 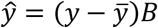. The transformations maximize the correlations between the pairs 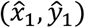 and 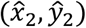 while ensuring the orthogonality of 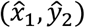 and 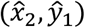, and equalize the variance of all canonical variables to one. The inclusion of cortical depth (*x*_3_) allowed us to compensate for slight departures of the objective from the surface normal.

The outcome of canonical correlation analysis is a representation of each bouton in the population by its canonical coordinates in cortical space 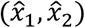 and its canonical coordinates in visual space (*ŷ*_1_, *ŷ*_2_). Using this representation of the data we can now compute the fluctuation of the density of ON/OFF cells in the canonical cortical domain, the fluctuation in the density of ON/OFF receptive fields centers in the canonical visual field, and test if they correlate with each other.

We used kernel density techniques to estimate the probability distribution of ON and OFF cells in canonical cortical space, denoted by 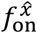 and 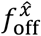 respectively (**Methods**). Similarly, we estimated the probability distribution of ON and OFF receptive fields centers in canonical visual space, yielding 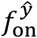 and 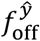. Fluctuations in the spatial distribution of ON and OFF cells in canonical cortical space, were calculated as the difference 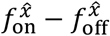. Similarly, to measure fluctuations in the distribution of ON and OFF receptive field centers in canonical visual space, we calculated the difference 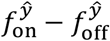 (**Fig. 3**).

**Fig 3.**
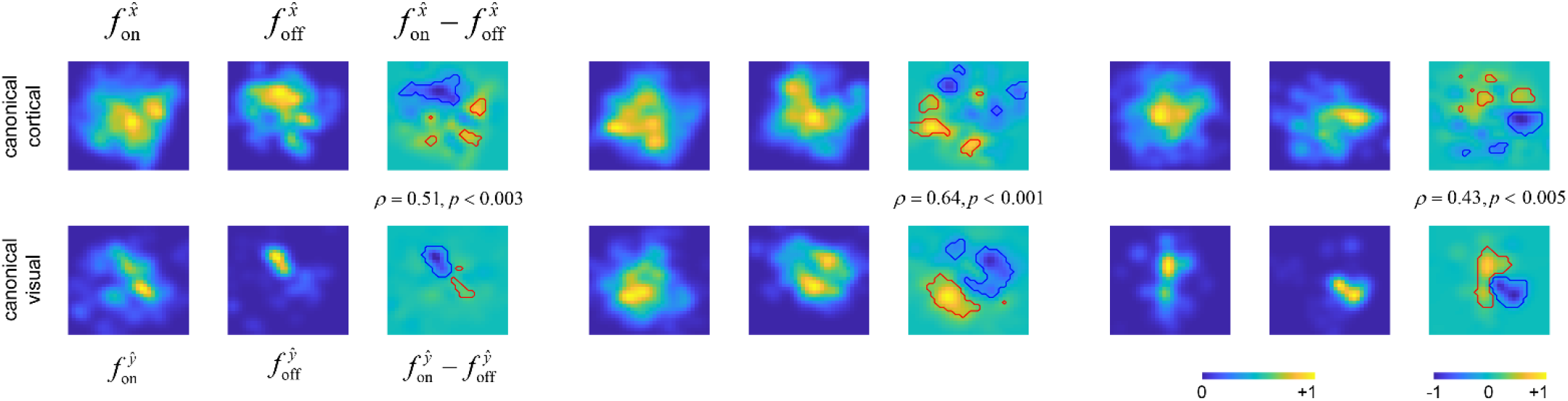
Fluctuations in the density of ON/OFF boutons correlates with the density of ON/OFF-center receptive field centers on the visual field. Each panel corresponds to a separate experiment. The top row of each panel shows, from left to right, the density of ON boutons in canonical cortical space, the density of OFF boutons, and their difference. The bottom row of each panel shows, from left to right, the density of ON-center receptive fields in canonical visual field, the density of OFF-center receptive fields, and their difference. We see that in all cases the fluctuations correlate. Densities are normalized to their peak. Fluctuations are normalized to their maximum absolute value.

The distributions of 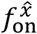 and 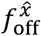 tend to be patchy, peaking in different locations, which results in the difference 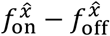 having statistically significant peaks and troughs (**Fig. 3**, top rows of each panel). We assessed the likelihood that the observed magnitudes in the fluctuations of 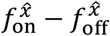 could arise by chance using Monte Carlo simulations where ON/OFF labels were randomly shuffled (**Methods**). Level sets were computed corresponding to the *p* = 0.001 significance level (**Fig. 3**, red and blue solid curves). As expected, we observe a clustering of ON/OFF boutons in the transformed canonical space as well (as the linear transformation is expected to preserve the structure we observed in native cortical space).

Similarly, we can calculate the density of receptive field centers for ON 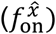 and OFF 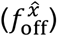 boutons, as well as their fluctuations, 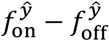 (Fig 3, bottom rows of each panel). These data corroborate our prediction – fluctuations in 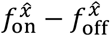 are mirrored by fluctuations in the balance of ON/OFF receptive field centers on canonical visual field, 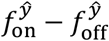. This is evidenced by the correlation between these functions (**Fig 3**, inset). Statistical significance was established by computing the likelihood that the observed level of correlation could arise by chance in controls that randomly shuffled ON/OFF labels (**Fig. 3**, *p-values*, **Methods**).

## Discussion

We have recently showed that, like other mammals (1–4), mouse primary visual cortex is parceled into ON/OFF domains (17). Those data revealed that fluctuations in the density of ON/OFF neurons on the cortical surface, which define ON/OFF domains, were mirrored by fluctuations in the density of ON/OFF receptive field centers on the visual field. We proposed that the input to the cortex itself might be biased in its representation of ON/OFF signals. The present study was designed to investigate this question by measuring the spatial distribution of ON/OFF thalamic boutons in the cortex.

We found that, as predicted, thalamic boutons cluster according to their responses to light increments and decrements, lending support to the so-called biased input model (17, 24). These findings are consistent with the notion that cortical receptive fields might be shaped by the spatial distribution of ON and OFF inputs in the visual field (11, 14, 18, 19). It would be important to conduct similar experiments in other mammalian species to ascertain if our results apply in general or are restricted to the mouse.

Finally, we note that we have not yet conclusively demonstrated that the imbalances observed in the thalamic input correlate with ON/OFF domains in individual animals. We are currently testing if this is the case by performing dual-color imaging where we reconstruct both the distribution of ON/OFF thalamic boutons and the ON/OFF responses of cortical neurons. We predict the fluctuations between ON/OFF signals in boutons and neurons will correlate.

## Acknowledgements

This work was supported by NIH grant NS116471 (D.L.R.)

## Contributions

E.T. performed all surgeries. D.L.R. conceived the study, collected/analyzed the data, and wrote the manuscript.

